# No allocentric distractor-location learning following self-movement even with contextual anchors and explicit knowledge

**DOI:** 10.64898/2026.11.16.767044

**Authors:** Litian Chen, Caterina Fatato, Shichen Jia, Freek van Ede, Heleen A. Slagter

## Abstract

Adaptive behavior requires learning where relevant information can likely be found, but also where distraction is likely to occur. For such learning to be useful in everyday life, it must generalize across changing viewpoints as observers move through the environment. Recent work suggests a striking asymmetry in this regard: whereas target-location learning can be grounded in world-centered coordinates, distractor-location learning appears tied to the observer’s current viewpoint. One possibility is that allocentric distractor-location learning requires stronger contextual support than allocentric target-location learning. Alternatively, distractor suppression may be fundamentally constrained by viewer-centered spatial coding. To distinguish between these possibilities, participants performed an additional-singleton search task displayed on a tabletop and continuously changed their standing position around the table, while the high-probability distractor location remained fixed in the external environment. Unlike previous studies using this paradigm, contextual anchors were embedded directly within the search display, allowing distractor locations to be encoded relative to a stable task-relevant allocentric structure. In a second experiment, participants were additionally explicitly informed about the distractor-location regularity. Despite task-relevant contextual structure and explicit knowledge, distractor-location learning failed to emerge in world-centered coordinates. Together with previous findings showing robust egocentric distractor-location learning using the same set-up, these findings strengthen the view that learned distractor suppression relies primarily on viewer-centered spatial coding, potentially reflecting a process linked to viewpoint-specific action selection.

## 1 Introduction

Adaptive behavior depends on the ability to learn from experience what in the environment is likely to matter and what can be safely ignored. Consistent with this idea, selective attention is not only shaped by what is currently goal-relevant or salient in the world, but also by selection history (Awh et al., 2012; Theeuwes et al., 2022). Through experience, observers’ attention becomes biased towards locations where behaviorally relevant stimuli are likely to occur (Anderson et al., 2021; Geng & Behrmann, 2005), while locations where salient distractors frequently occur are increasingly suppressed or ignored (Theeuwes, 2025; van Moorselaar & Slagter, 2020; Wang & Theeuwes, 2018). These attentional biases develop rapidly and are generally assumed to reflect implicit learning of statistical regularities in the environment.

When agents actively move through the environment, however, spatial regularities can be represented either relative to the agents’ current viewpoint (*egocentric* reference frame) or relative to stable locations in the external world (*allocentric* reference frame; Colby, 1998). Recent studies suggest a striking difference between target- and distractor-location learning in this respect. Using a dynamic visual-search task in which participants continuously changed their standing position around a tabletop display, we previously showed that that implicit distractor-location learning emerges when the likely distractor location is defined in egocentric, viewpoint-specific coordinates, but not when it is stable in the external world instead (Chen, van Ede, Jungerius, & Slagter, 2026). Critically, even explicitly informing participants about the world-centered high-probability distractor location could not yield allocentric distractor-location learning (Chen, van Ede, & Slagter, 2026). This contrasts with target-location learning, where explicit knowledge can enable observers to exploit allocentric spatial regularities across changes in viewpoint (Chen, van Ede, & Slagter, 2026; Jiang et al., 2013; Jiang & Swallow, 2014).

These findings suggest that while *target*-location learning may generalize across viewpoints to support attentional guidance in dynamic environments, learned *distractor*-location learning may be inherently constrained by egocentric spatial coding. This distinction may arise because targets must be selected to guide ongoing behavior, making it useful to represent their likely locations across changing viewpoints, while distractors matter only insofar as they interfere with ongoing behavior. The task therefore may not be to encode where they are in the world, but to regulate their influence on attentional selection, when encountered. Our previous findings showing no allocentric distractor learning despite explicit knowledge (Chen, van Ede, & Slagter, 2026) suggest that this interference is intrinsically viewer-centered: distractors compete with targets for selection within the observer’s current field of view. If learned suppression is primarily implemented at this level, explicit knowledge of distractor locations may provide limited benefit, as such knowledge may not readily translate into more effective proactive control over distractor interference. Consistent with this possibility, distractor suppression appears largely insensitive to explicit intentions and top-down knowledge. Explicitly cueing a distractor location often fails to reduce distractor interference (Addleman & Störmer, 2023; Becker et al., 2015; Cunningham & Egeth, 2016; Wang & Theeuwes, 2018), and explicit knowledge of a distractor-location regularity does not enhance distractor-location learning (Gao & Theeuwes, 2022). By contrast, target-location cueing typically facilitates attentional selection and improves performance (Posner, 1980; Wolfe, 2021). Together, these findings raise the possibility that distractor suppression is implemented within viewer-centered priority maps, constraining learning to the observer’s current perspective.

An alternative possibility is that the absence of allocentric distractor-location learning, even with explicit knowledge, reflects insufficient task-relevant spatial context rather than an inherent constraint on learned distractor suppression. Although participants had access to stable environmental cues, including room landmarks (e.g., a window) and a fixed marker on one side of the computer display, these cues were external to the search task itself and therefore not part of the task-relevant spatial configuration that participants actively used during search. Moreover, as is typical in additional-singleton visual-search tasks, the search items were arranged in a circular layout, rendering the search configuration visually identical across viewpoints. Thus, although stable allocentric cues were available in the environment, the search task itself provided little intrinsic spatial structure for anchoring the likely distractor location across changes in viewpoint. As a result, explicit knowledge may have failed to support allocentric distractor-location learning because participants lacked task-relevant spatial anchors for establishing a stable allocentric representation of the likely distractor location.

This possibility is supported by recent work in naturalistic virtual environments showing that stable anchor objects embedded in the relevant environment can support allocentric coding of nearby object locations, particularly when these anchors are proximal to the encoded location (Baltaretu et al., 2024, 2026). Likewise, the use of surrounding objects as allocentric cues depends on their reliability, spatial configuration, and proximity to the encoded location (Klinghammer et al., 2017). In the context of visual search, embedding additional contextual anchors directly within the search display may therefore provide the relational information needed to support world-centered distractor-location learning across changes in viewpoint. Such a local structure may be particularly important when observers can exploit explicit knowledge of a likely distractor regularity to translate it into a stable, perceptually grounded, allocentric distractor-location representation.

Nevertheless, the absence of allocentric distractor-location learning is unlikely to be explained solely by the inability of participants to notice the contextual structure. Using the exact same circular search display, task, and setup, we previously found that explicit knowledge enabled allocentric target-location learning (Chen, van Ede, & Slagter, 2026). Thus, participants evidently had sufficient access to stable environmental information to support allocentric learning when the learning concerned likely target locations. This suggests that either allocentric distractor-location learning requires richer or more task-relevant contextual structure or that distractor suppression and target facilitation rely on distinct spatial-learning mechanisms.

The present study was designed to adjudicate between these two possibilities and determine whether allocentric distractor learning *can* emerge when the environment provides richer, task-relevant contextual structure that is embedded within the configuration of the task-relevant search display itself. In two experiments, participants performed an additional-singleton search task while continuously changing their standing position around a tabletop search display. As in our previous study, the high-probability distractor location remained fixed in the external environment. Critically, however, we embedded the anchors, e.g. the two additional search items placed at fixed positions near one border of the display, directly within the search display itself as the task-relevant contextual structure (Figure 1). These anchors formed part of the task-relevant search environment and created a stable spatial configuration that, unlike the previously used circular display, no longer appeared identical across viewpoints. As such, the search display provided a continuously available and proximal reference structure for representing locations. In Experiment 1, participants were not informed of the distractor-location regularity, allowing us to test whether such contextual structure is sufficient to support incidental allocentric distractor-location learning. In Experiment 2, participants were explicitly informed of the high-probability distractor location, allowing us to examine whether with explicit knowledge, this enriched contextual structure could be exploited to induce allocentric distractor-location learning. We reasoned that if previous failures to observe allocentric distractor-location learning resulted from insufficient task-relevant contextual support, then embedding stable contextual anchors within the search display should enable allocentric learning. In this case, distractor interference should be reduced when distractors appear at the high-probability allocentric location in both Experiment 1 and 2 (just as we and others have repeatedly and robustly observed for egocentric distractor-location learning when keeping the high-probability distractor location fixed from the viewpoint of the participant; Chen, van Ede, Jungerius, & Slagter, 2026; Chen, van Ede, & Slagter, 2026; Jiang et al., 2014; Jiang & Swallow, 2013). In contrast, if distractor suppression is fundamentally constrained by viewer-centered coding, enriching the allocentric context and even providing explicit knowledge should not overcome this limitation.

**Figure 1.**
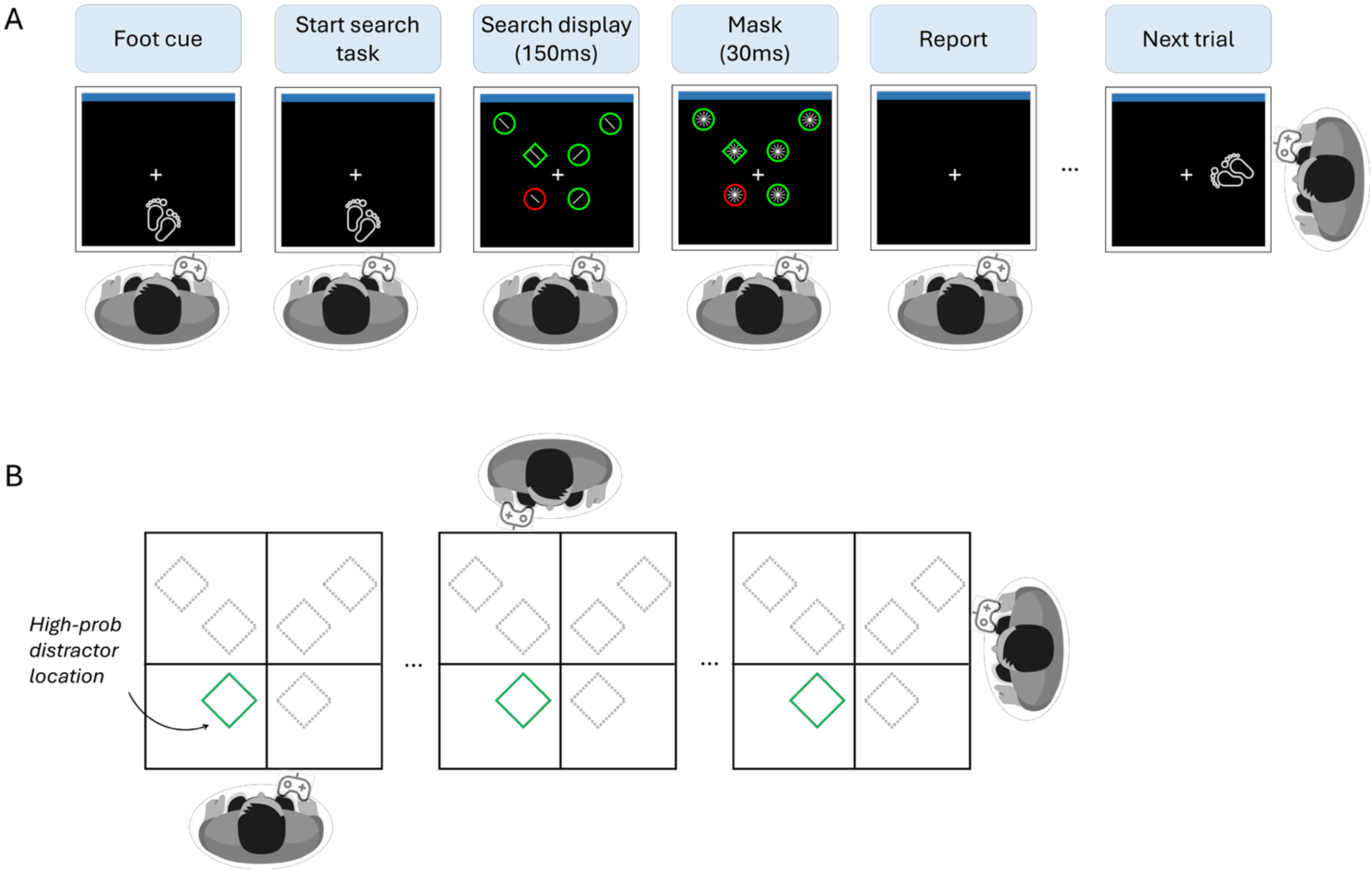
(A) Schematic illustration of a task trial. At the beginning of each trial, a foot cue indicated the side of the table at which the participant should stand. After moving to the indicated position, participants simultaneously pressed two keys on a handheld button box to initiate the search task. The search display was presented for 150 ms, followed by a 30-ms mask. Participants had to indicate whether the line element inside the target (the unique shape) was tilted to the left or right and ignore the singleton distractor (the unique color). Targets and singleton distractors were presented at all six locations. Their required standing position varied across trials, resulting in continuous changes in viewpoint throughout the experiment. (B) Spatial schematic illustrating that the high-probability distractor location was stable in the environment. Independent of the participant’s standing position, it always appeared more often in the same location (search quadrant).

## Experiment 1

### 2.1 Methods

The aim of Experiment 1 was to determine whether stable contextual anchors embedded within the search display itself can support allocentric distractor-location learning in the absence of explicit knowledge.

### 2.2 Participants

Twenty-four participants participated in Experiment 1 (22 female, *Mean age* = 20.0, *SD* =1.9). The sample size was determined a-priori using a simulation-based power analysis with the effect size of the egocentric distractor-location learning effect observed in our previous study (https://osf.io/7g8a4/) (Chen, van Ede, Jungerius, & Slagter, 2026). The study was approved by the Scientific and Ethical Review Board of the Vrije Universiteit Amsterdam. Participants were drawn from a pool of university students that either participated for course credits or money (12.5 euros/hour). Written informed consent was collected at the onset of the study.

### 2.3 Apparatus and stimuli

The task was coded in MATLAB 2019a (www.mathworks.com) and adapted from Chen, van Ede, & Slagter (2026). It is depicted in Figure 1. Participants performed the task while standing right next to a square table (length x width x height: 80 x 80 x75 cm). The surface of the table was made of high-quality semi-transparent optical glass that minimized light refraction from all angles of the table. The task contents were projected on this surface using a beamer positioned underneath the table. Participants used a wireless numeric keypad to respond.

### 2.4 Procedure and design

As shown in Figure 1, participants performed an additional-singleton visual search task in which they had to search for a unique shape (the target) among five non-target shapes and indicate whether the line inside the target shape was tilted left or right. They continuously changed their standing position, and hence, their viewpoint, from trial-to-trial. In 66.7% of trials, a singleton distractor (shape with a unique color) was present. In 66.7% of such distractor-present trials, the singleton occurred more often at one of the locations. Critically, this high-probability distractor location was fixed in the world (i.e., viewpoint independent).

Each search display consisted of six shapes. All stimuli were presented on a black background (RGB values = 0, 0, 0). Each shape was drawn in a red (RGB values = 255, 0, 0) or green (RGB values = 0, 255, 0) outline with a 45° tilted white line segment (1.6°) at their center. The six shapes included one circle (radius 1.05°) and five diamonds (2.7° × 2.7°), or vice versa. Targets could occur at each shape location with equal probability. To make sure that stimuli were comparable in size across participants, the size of stimuli in pixels was adjusted based on the individual height of each participant. In our previous studies (Chen, van Ede, & Slagter, 2026), the search elements were arranged in a circular configuration, so that the search configuration appeared identical across viewpoints, which may have prevented allocentric coding. In contrast, in the current work, while four of the shapes were equally spaced along an imaginary circle (radius = 4.3°) centered on the fixation cross, the other two shapes were presented outside the imaginary circle at fixed locations symmetrically positioned near one of the four borders of the search display. Crucially, these additional shapes (the anchors) formed part of the search display itself and therefore created a stable task-relevant contextual structure that looked different across viewpoints. The locations of the anchors had four possible configurations (top, bottom, left, or right border) and were randomly assigned and counterbalanced across participants. In addition, as in our previous study (Chen, van Ede, Jungerius, & Slagter, 2026), one thick blue bar (RGB: [46 117 182]) was shown continuously at the same physical location, on one side of the computer display, and served as another local environmental landmark.

At the beginning of each trial, a foot picture randomly shown on one of the four sides of the display indicated at which side of the table the participant should stand (and move to) for that trial. Once in the correct position, participants were instructed to focus on the fixation cross and press the two response buttons (key1 and key3) simultaneously to start the search task. After 50 ms, a search display was then presented for 150 ms, followed immediately by a mask for 30 ms. The mask consisted of a set of six white line segments which were placed at the center of each stimulus (cf. van Moorselaar & Theeuwes, 2021). Participants were instructed to indicate whether the line inside the target was tilted to the left or right by pressing the corresponding response button within 3000 ms. They were asked to respond as quickly and accurately as possible. Sound feedback was provided if participants made a wrong report or if the allowed response time (3000 ms) was up.

The task was divided into two parts: a learning phase and an extinction phase. The learning phase consisted of eight blocks of 90 trials. During both phases, a color-singleton distractor was present on two thirds of the trials (distractor-present trials, 60 trials), whereas all six stimuli had the same color on the remaining one third of trials (distractor-absent trials, 30 trials). Within each learning block, when the singleton distractor was present, it occurred more often (in 66.7% of distractor-present trials, in total 40 trials) at one of the four locations around fixation (high-probability distractor location; randomized across participants). The singleton occurred equally often at the other five low-probability locations on the remaining distractor-present trials (20 trials). Targets were distributed equally across the six possible locations in both distractor-present and distractor-absent trials. The subsequent extinction phase consisted of four blocks with 90 trials each, during which the spatial distractor regularity introduced during the learning phase was removed. That is, both distractors and targets were now presented equally often at all six possible locations.

In Experiment 1, participants were not explicitly informed about the spatial distractor-location regularity. To test if incidental awareness of the regularity may have developed as a function of learning, at the end of the task, participants completed an awareness test. Participants were asked to choose one location on the search display that they felt the uniquely colored item may have occurred more often in the search task, as well as their confidence in that choice (scaled from 1 to 4, ranging from 1: completely guessing, 2: somewhat confident, 3: almost certain, to 4: completely certain).

### 2.5 Data analysis

All data analyses were conducted in R. Following our previous work (Chen, van Ede, Jungerius, & Slagter, 2026; Chen, van Ede, & Slagter, 2026), we first determined whether the mean reaction times (RTs) for each participant were above or below 2.5 standard deviations from the group mean RTs to remove participants with super-slow or fast responses. No participants were excluded using this criterion. We then applied a two-step trimming procedure. First, incorrect trials (18.8 %) and trials with reaction times shorter than 200 msec (2.01 %) were excluded from the data set. Then, trials were trimmed based on a cutoff value of 2.5 standard deviations from the mean RT per participant (2.98 % trials removed).

We next examined the presence of distractor-location learning using paired t-tests contrasting mean reaction times between high and low distractor-probability location conditions in the learning phase. Because capture is reduced for distractors further away from fixation, trials in which either the distractor or the target occurred at the anchors locations were excluded from this and all other analyses for both distractor-present and distractor absent trials (the inclusion of these two additional locations exclusively served to provide a contextual allocentric anchor). If the task-relevant contextual structure supported allocentric distractor-location learning, reaction times were expected to differ between the high- and low-probability conditions. In contrast, the absence of search time differences would indicate no evidence for allocentric distractor-location learning.

We then examined whether any distractor-location learning observed in the learning phase would remain in the extinction phase, after removal of the spatial distractor regularity. We previously found that egocentric distractor location learning quickly disappeared and was only still present in the first extinction block (Chen, van Ede, Jungerius, & Slagter, 2026). Therefore, we conducted a repeated measures ANOVA contrasting reaction times between trials where the distractor appeared at what used to be the high probability location versus trials where it occurred at any of the other 3 central, previously low probability locations to check for any remnants of learning, with Distractor Location and Block as within-subject factors. It should be noted that the extinction phase yielded only 5.2 trials per participant and block on average for the high-probability condition (range: 2–8) and 14.6 trials for the low-probability condition (range: 9–19). Therefore, we decided not to explore potential effects of distractor location learning per block given these low trial numbers.

For the final awareness task, we used a binomial test to determine whether the proportion of participants who correctly identified the high-probability distractor location exceeded chance level (i.e., 1/6, given the six possible locations), thereby serving as a group-level index of incidental awareness.

### 2.6 Results

#### 2.6.1 Distractor-location learning in the learning phase

As shown in Figure 2, distractors generally impaired performance, as reflected in slower response times in trials in which the distractor appeared (at either a low-probability or a high-probability location) compared to the distractor-absent trials. However, no evidence for distractor-location learning was found: RTs were comparable for high- and low-probability distractor location tirals (high vs. low: 769 ms vs. 770 ms, MD = 0.74 ms, 95% CI [−18.65, 20.13], *t*(23) = 0.08, *p* = .938, BF_01_ = 4.65). Thus, when the distractor-location regularity was fixed in the world, we did not observe allocentric distractor-location learning, even with additional task-relevant contextual structure. This strongly contrasts with the learning effects we previously reported (Chen, van Ede, Jungerius, & Slagter, 2026) and replicated (Chen, van Ede, & Slagter, 2026) when keeping the regularity fixed in a view-point dependent manner.

**Figure 2.**
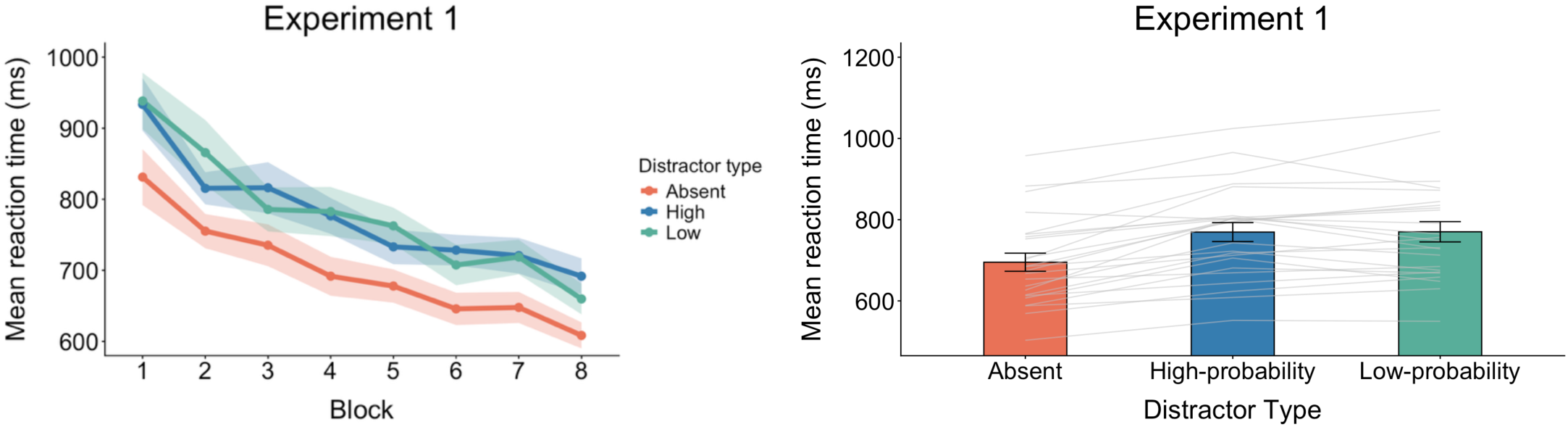
Results of Experiment 1 investigating implicit allocentric distractor-location learning. A) Search times over time in the learning phase, as a function of block and trial type. As can be seen, search times were equally slowed down by singleton distractors at high-versus low-probability locations, suggesting an absence of allocentric distractor-location learning. B) Mean search time across blocks per trial type in the learning phase, with individual participant data shown with thin grey lines. Error bars in all plots depict ± 1 SEM

#### 2.6.2 Extinction phase

Although no distractor-location learning was observed in the learning phase, we still conducted our planned analysis exploring whether any distractor-location learning persisted after the regularity was removed in the extinction phase. As expected, given the absence of learning in the learning phase, the repeated measures ANOVA results showed no overall effect of distractor-location learning in the extinction phase as reflected in the absence of a significant main effect of Distractor Location: *F*(1, 23) = 1.68, *p* = .208, *η*²*_p_* = .068, and a non-significant interaction between Distractor Location and Extinction Block (*F*(2.80, 64.34) = 0.40, *p* = .736, *η*²*_p_* = .017).

#### 2.6.3 Awareness test

Finally, we statistically examined incidental awareness of the distractor regularity. Eight participants indicated the correct location, while sixteen participants chose an incorrect location. A one-sided exact binomial test showed that this percentage (33.3%) was significantly above the 1/6 chance level, *p* = .035. Of the eight participants who reported the correct location, six reported being somewhat confident (confidence level ‘2-somewhat’) in their choice, two reported that they were guessing (confidence level ‘1 - Guessing’) and none reported being almost or completely certain of their choice. Thus, although there was some evidence of above-chance knowledge of the high-probability location at the group level, confidence in these responses was generally low.

To explore if potential awareness had affected distractor-location learning, we conducted an additional analysis comparing the effect of the distractor regularity on search times between participants who successfully selected the likely singleton distractor location (allegedly aware, though guessing correctly also remains a possibility here) and who did not (unaware). As can be seen in Figure 3, results showed no significant difference in this effect between “aware” and unaware participants: *t*(22) = 0.77, *p* = .448, Cohen’s *d* = 0.33. There was no evidence for allocentric distractor learning in either group (aware group: M = 11.06 ms, *t*(7) = 0.72, *p* = .497, Cohen’s *d* = 0.25; unaware group: *t*(15) = -0.37, *p* = .715, Cohen’s *d* = -0.09).

**Figure 3.**
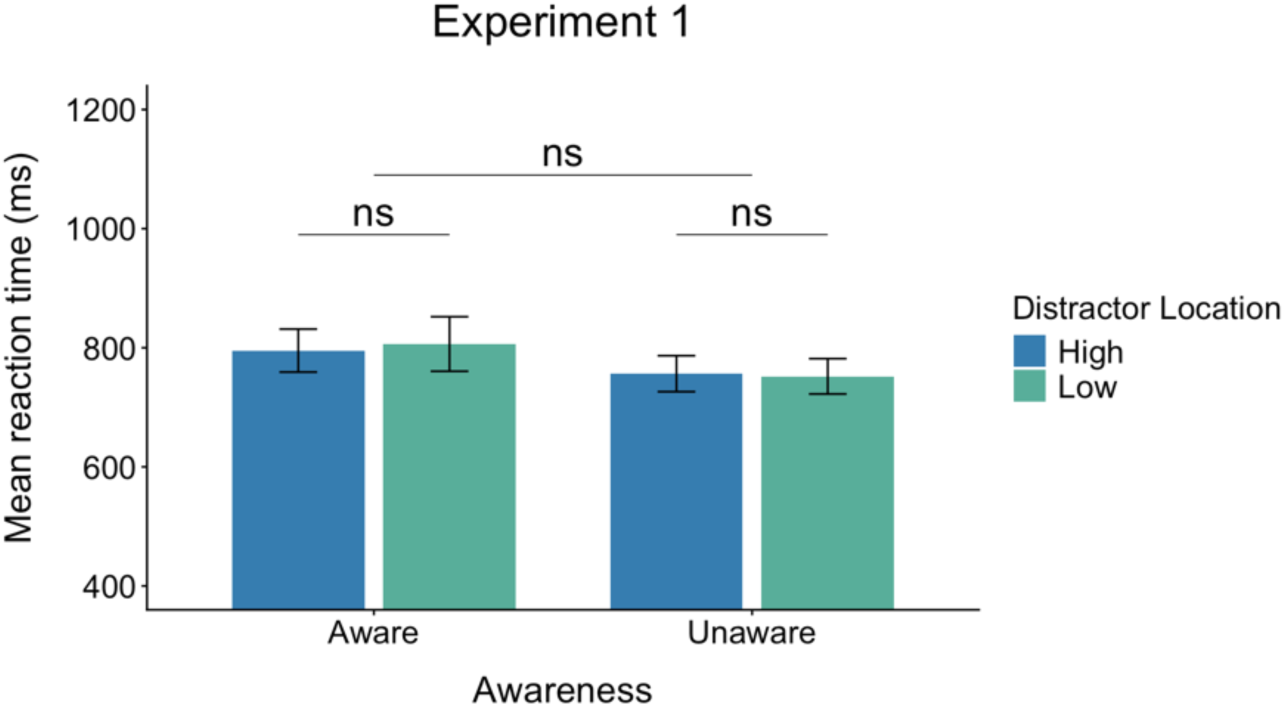
No effect of post-hoc correct identification of the likely distractor location on distractor-location learning in Experiment 1. Participants who correctly (’Aware’) reported the high probable distractor location during the Awareness test and those who were incorrect (’Unaware’) both displayed no significant difference in search times between trials in which the distractor occurred at the high-probable versus low-probable location. *n.s.* = not significant. Error bars in all plots depict ±1 SEM.

### 2.7 Discussion

In Experiment 1, we did not observe that embedding stable contextual anchors within the search display induced reliable allocentric distractor-location learning. This replicates and extends past findings (Chen, van Ede, Jungerius, & Slagter, 2026; Chen, van Ede, & Slagter, 2026) by showing that allocentric distractor-location learning does not emerge *even* in the presence of task-intrinsic spatial structure. This failure to observe allocentric distractor-location learning is notable as we previously found robust egocentric distractor-location learning using the same paradigm (although without task-relevant allocentric structure) (Chen, van Ede, Jungerius, & Slagter, 2026; Chen, van Ede, & Slagter, 2026) and given a large body of work that has consistently shown rapid and robust distractor-location learning in static search environments (Britton & Anderson, 2020; Kim & Anderson, 2022; Theeuwes et al., 2022; van Moorselaar et al., 2021). Although we did not observe any difference between participants who correctly identified the high-probability distractor location and those who did not, distractor-location knowledge was assessed only at the end of the experiment. It therefore remains unclear when this knowledge emerged and whether it was available early enough to influence learning. Moreover, participants were generally not very confident in their location choice, even when correct, and may therefore have not exploited the knowledge. Finally, our measure does not clarify whether a correct location choice reflects explicit knowledge, implicit knowledge, or mere guessing. It therefore remains possible that allocentric distractor-location learning may emerge in the presence of task-intrinsic spatial structure when also having explicit knowledge of the distractor regularity from the outset. Experiment 2 tested this possibility.

### Experiment 2

Experiment 1 examined whether embedding additional contextual anchors directly within the search display could provide sufficient spatial context to enable implicit allocentric distractor-location learning. However, no reliable distractor-location learning was observed. Previous work on target-location learning (Jiang et al., 2013) has shown that explicit knowledge can promote allocentric learning, enabling observers to exploit world-centered spatial regularities that are not expressed under implicit learning conditions. More recently, we replicated and extended that finding showing that explicit knowledge facilitated allocentric target-location learning, but failed to induce allocentric distractor-location learning (Chen, van Ede, & Slagter, 2026). Notably, however, this earlier work employed a circular search display that provided little intrinsic spatial structure across viewpoints. Therefore, in Experiment 2, we tested whether explicit knowledge of the distractor-location regularity combined with the contextual anchors we introduced in Experiment 1 would enable allocentric distractor learning.

### 3.1. Participants

Twenty-four participants participated in Experiment 2 (17 female, Mean age = 19.8, SD =1.8). Two participants were excluded from analyses because in the post-hoc awareness test, they selected the wrong location as the high-probability distractor location, potentially suggesting that they may have not been well aware of the high-probability distractor location. As in Experiment 1, participants were drawn from a pool of university students that either participated for course credits or money (12.5 euros/hour). Written informed consent was collected beforehand. The study was approved by the Scientific and Ethical Review Board of the Vrije Universiteit Amsterdam.

### 3.2. Procedures and design

The task and procedure in Experiment 2 were identical to the task and procedure in Experiment 1, except as indicated in the following. First, participants were explicitly informed of the high-probability distractor location at the onset of the experiment, after practicing the task first. Specifically, they were told that the singleton distractor was more likely to appear at one particular location, defined as a fixed location on the monitor (allocentric), independent of their position. The instruction was as follows: “*As you probably noticed in the practice, on some trials one of the non-target shapes had a unique colour (e.g. red among all green shapes or vice versa). Here, the uniquely colored (green) distractor is shown at the current location (high-probability location) where it will occur on the monitor on 66.7% of trials when it is present. This means this distractor occurs much less often on the other monitor locations. The high-probability distractor location is fixed on the monitor, no matter where you stand. While the target (the unique shape) occurs at each monitor location equally often. Remember, you can ignore this distractor! It is not relevant to the task. Remember, the high-probability distractor location is fixed on the monitor. So, when you walk around the table, this does not change its likely location on the monitor. In other words, the likely distractor location is not dependent on your standing position.”*

Next, to ensure that participants had acquired explicit knowledge of the high-probability distractor location, they needed to successfully indicate the high-probability distractor location by pressing the button that corresponded to the number shown at that location, from three standing positions. Only once they passed these three location tests, they could continue with the formal task. During the learning phase, at the start of each block starting from block 2, participants again had to first pass the location test from three different standing positions to ensure that they remembered the high-probability distractor location. If participants selected an incorrect location, they received sound feedback, and the task proceeded to the next trial. At the end of each location test, participants were given another opportunity to review the high-probability distractor location.

At the end of the task, participants completed a final awareness test. They again performed the location test, indicating the high-probability distractor location from three different standing positions, and their selections were collected immediately without any feedback. They additionally completed a short questionnaire that included rating the extent to which they used the explicit knowledge about the distractor location to ignore the distractor and how helpful they found this knowledge for ignoring the distractor on a scale from 1-7. These two questions were formulated as follows: “*Did you use the knowledge about the likely distractor location to ignore it? Press the button between 1 to 7 to give a rank of your feeling*”, “*Did the knowledge about the likely distractor location help you ignore it? Press the button between 1 to 7 to give a rank of your feeling*”.

### 3.3. Analysis

Data cleaning followed the same procedure as Experiment 1. In total, 18.40% of trials were removed as incorrect responses, 1.04% as anticipatory responses (RT < 200 ms), and a further 3.07% by the ±2.5 SD RT trimming criterion.

We again first examined the presence of allocentric distractor-location learning using a paired t-test contrasting mean reaction time between high- and low-probability distractor location trials in the learning phase (1-8 blocks). This analysis was also restricted to trials in which both the target and the singleton distractor appeared at one of the four central locations.

In the extinction phase, based on the data from Experiment 1 and previous work, we expected no overall learning effect of the previous high-probability distractor location. To verify this, we conducted a repeated measures ANOVA with Distractor Location (previously high, previously low) and Block as within-subject factors. As in Experiment 1, trial numbers per block were low: the extinction phase yielded only 4.8 trials per participant and block on average in the high-probability condition (range: 2–8) and 13.6 in the low-probability condition (range: 7–19). Therefore, we did not conduct any comparisons per block separately.

For the awareness task, we checked that participants selected the correct high-probability distractor location in the three test trials at the end of the experiment. Participants who did not were excluded from the analyses (n=2).

### 3.4 Results

#### 3.4.1 Distractor-location learning in learning phase

As shown in Figure 4, distractors generally impaired performance, as reflected in slower response times when a distractor was present compared to distractor-absent trials. However, again we obtained no evidence for allocentric distractor-location learning, even with additional contextual anchors as well as explicit knowledge. RTs were comparable between trials in which the distractor occurred at the high-versus low-probability locations (high vs. low: 756 ms vs. 765 ms, MD = 9.32 ms, 95% CI [−9.22, 27.87], *t*(21) = 1.05, *p* = .308, BF_01_ = 2.77).

**Figure 4.**
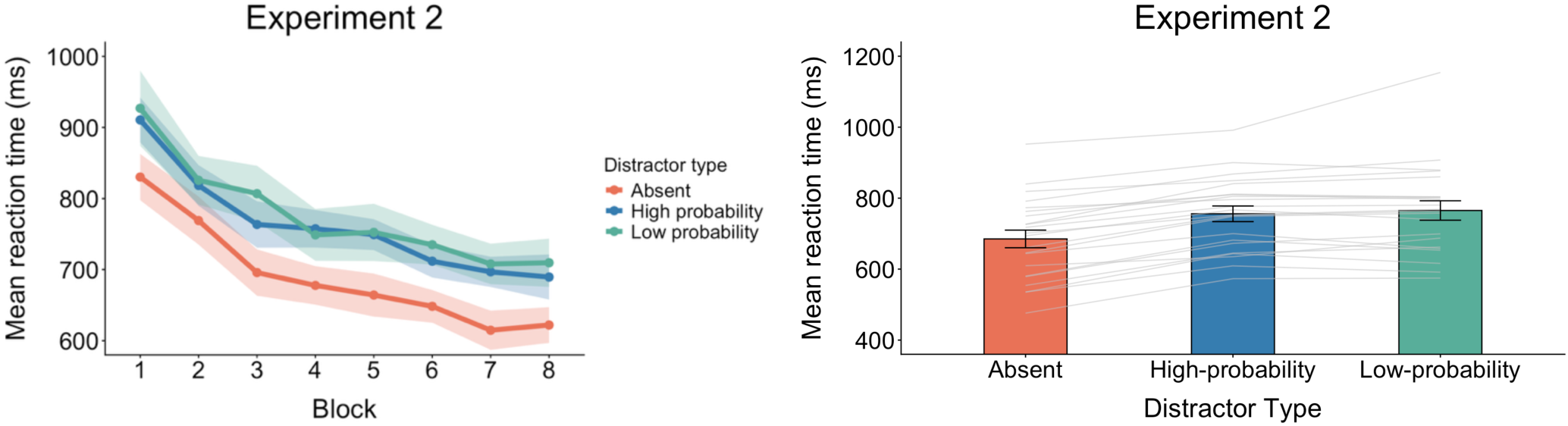
Results of Experiment 2 investigating explicit allocentric distractor-location learning. A) Search times over time, as a function of block and trial type in the learning phase. As can be seen, explicit knowledge did not induce distractor location learning as search times did not differ between high- and low-probability trials over time. B) Mean search times across blocks per trial type in the learning phase. Error bars in all plots depict ± 1 SEM.

#### 3.4.2 Extinction phase

Although the learning phase provided no evidence for allocentric distractor-location learning with explicit knowledge, we nonetheless explored whether effects of allocentric distractor-location learning were detectable after the spatial regularity was removed in the extinction phase. The repeated measures ANOVA results showed no remnants of distractor-location learning in the extinction phase, as reflected by a non-significant main effect of Distractor Location: *F*(1, 21) = 0.58, *p* = .456, *η*²*_p_* = .027, and a non-significant interaction between Distractor Location and Extinction Block (*F*(2.63, 55.27) = 0.53, *p* = .638, *η*²*_p_* = .025).

#### 3.4.3 Awareness test

In the final location awareness test, 22 participants selected the correct location all three times, suggesting that most participants had explicit knowledge of the high probability distractor location despite showing no evidence for allocentric distractor location learning. Participants’ answers to the question whether they used the distractor location knowledge to ignore the distractor suggested low use on average (*M* = 2.45, *SD* = 1.14, 95% CI [1.95, 2.96]). They furthermore rated the perceived helpfulness of this knowledge relatively low (*M* = 3.27, *SD* = 1.58, 95% CI [2.57, 3.97]).

### 3.5 Discussion

In Experiment 2, even when participants were explicitly informed of the distractor regularity in our enriched allocentric stimulus configuration from the outset, we did not observe reliable allocentric distractor-location learning. Explicit knowledge therefore did not promote allocentric coding or learning of the distractor location despite the presence of an enriched allocentric context.

## General discussion

The central question addressed in the present study was whether distractor-location learning can be anchored to stable locations in the external world when observers continuously change their viewpoint. Across two experiments, we found no evidence for allocentric distractor-location learning despite the availability of external environmental landmarks, a stable task-relevant spatial structure embedded within the search display, and, in Experiment 2, explicit knowledge of the world-centered high-probability distractor location. This failure to observe allocentric distractor-location learning is notable as we previously found robust egocentric distractor-location learning using the same paradigm (although without task-relevant allocentric structure) (Chen, van Ede, Jungerius, & Slagter, 2026; Chen, van Ede, & Slagter, 2026) and given a large body of research consistently showing rapid and robust distractor-location learning in static search environments (Addleman & Störmer, 2023; Awh et al., 2012; Britton & Anderson, 2020; Gaspelin & Luck, 2018; Kim & Anderson, 2022; Theeuwes et al., 2022; van Moorselaar et al., 2021). Together, these findings suggest that learned distractor suppression is strongly tied to the observer’s current perspective and constrained by egocentric spatial coding.

Importantly, the current results should not be taken to imply that allocentric distractor suppression is categorically impossible. Rather, they point to a more specific limitation: neither additional contextual structure nor explicit knowledge was sufficient to induce robust allocentric distractor-location learning while observers continuously changed their viewpoint through full-body movement. At first sight, this finding may appear surprising given that recent studies have demonstrated that distractor suppression can be modulated by contextual information. Observers can learn associations between visual or task contexts and distractor probabilities and such learning reduces distractor interference (Allon & Leber, 2025; Moher & Leber, 2025). However, the function of context fundamentally differed from that in the present studies. In those studies, contextual cues signaled the likelihood of distraction or the current task state, but they did not provide a stable spatial map for representing distractor locations, nor did they require observers to update spatial representations across changing viewpoints. In contrast, the present study embedded contextual information directly into the spatial structure of the search environment, such that it could serve as an allocentric reference frame for learning where distractors were most likely to occur. Moreover, participants continuously changed their viewpoint through full body movement, allowing us to examine world-based distractor learning. Despite these conditions, no allocentric distractor location learning established.

More broadly, the present findings help clarify the conditions under which learned distractor suppression can be expressed in different spatial reference frames. Recent studies have shown that suppression is not exclusively retinotopic and can under some circumstances be expressed in object-centered or spatiotopic coordinates (Chang & Golomb, 2025; Ilksoy et al., 2025a; Ilksoy et al., 2025b). Importantly, however, these studies primarily manipulated eye position or object structure while observers remained stationary. Thus, the relevant reference frames could still have been anchored to the observer. The present study applied a stronger test by asking whether distractor suppression can be tied to stable locations in the external world while observers continuously changed body position and viewpoint by walking around a tabletop. Despite the presence of task-relevant contextual structure and environmentally stable landmarks, and, in Experiment 2, explicit distractor knowledge, we found no evidence for allocentric distractor-location learning.

Experiment 2 further addresses the role of explicit knowledge in distractor-location learning. In line with our previous work (Chen, van Ede, & Slagter, 2026), the present results indicate that having explicit knowledge of a distractor regularity might not equate to being able to use this knowledge or implement it as a top-down attentional control strategy. This observation is consistent with previous studies showing that awareness of a distractor-location regularity does not necessarily strengthen distractor suppression (Gao & Theeuwes, 2022) and that explicit distractor cues are often ineffective in top-down reducing distraction (Addleman & Störmer, 2023; Noonan et al., 2016). Thus, the critical limitation may lie in translating the explicit knowledge of the distractor regularity into top-down spatial attentional suppression. Consistent with this, participants in Experiment 2 reported relatively low levels of strategically using their explicit knowledge of the high-probability distractor location. In Experiment 1, at the group level, participants identified the high-probability distractor location above chance in the post-experiment awareness test, but distractor-location learning was similarly absent in ’aware’ and ’unaware’ participants. This too suggests that even if some information about the spatial regularity is acquired, this does not facilitate allocentric distractor-location learning.

Interestingly, these findings contrast with evidence showing that explicit knowledge can enhance allocentric target-location learning under highly similar conditions (Chen, van Ede, & Slagter, 2026; Jiang et al., 2013; Jiang & Swallow, 2014). Although the present study did not directly compare target and distractor learning, our prior work revealed such a dissociation, and the present findings extend that pattern. Across multiple studies, allocentric target-location learning with explicit knowledge has proven robust, whereas allocentric distractor-location learning has consistently failed to emerge, even when observers possess explicit knowledge of the regularity. This pattern raises the possibility that attentional facilitation and suppression rely on partly distinct mechanisms, in line with previous proposals (Jensen, 2024; Noonan et al., 2016; van Moorselaar & Slagter, 2020). Whereas explicit spatial knowledge may readily support the anticipation of likely target locations and viewpoint-independent target-location learning, distractor suppression appears considerably less amenable to such top-down control and more constrained by egocentric coding.

One interpretation of the present findings is that distractor suppression is implemented through mechanisms that are closely linked to action control (Chen et al., 2026). When a distractor is encountered, suppressing its processing necessarily requires specifying its location relative to the observer. In other words, the implementation of distractor suppression depends on information represented in egocentric coordinates. Under this view, suppression does not primarily encode *where* distractors likely occur in the external world, but *how* attentional deployment should be suppressed relative to one’s current position and orientation. Such a mechanism would naturally favor viewer-centered learning and could explain why allocentric distractor-location learning repeatedly fails to emerge, even when observers possess explicit knowledge of the regularity and have access to stable contextual information.

The present findings do not rule out the possibility that even richer or more naturalistic environments may facilitate allocentric distractor-location learning. For example, meaningful anchor objects can support allocentric coding of nearby locations in naturalistic settings (Baltaretu et al., 2024, 2026). In general, natural scenes provide rich semantic and spatial structure that guides attention (Võ et al., 2019). Future studies could therefore examine whether richer and more naturalistic environmental landmarks facilitate allocentric distractor-location learning during active movement.

Taken together, the present findings suggest that neither explicit knowledge of a distractor-location regularity nor the availability of a stable task-relevant contextual structure is sufficient to establish allocentric distractor-location learning when moving through a three-dimensional environment. Together with prior findings, this suggests that learned distractor suppression, unlike learned target facilitation, is strongly grounded in the observer’s current perspective. These findings place important constraints on the extent to which contextual information and explicit spatial knowledge can shape distractor suppression across changing viewpoints. They suggest that learning to suppress distractors may be more closely linked to action-oriented control mechanisms operating in egocentric coordinates than in abstract world-centered spatial representations.

## Acknowledgements

This work was supported by an ERC Consolidator grant (PlasticityOfMind; 101002584) to H.A.S. F.v.E. was supported by an NWO Vidi Grant from the Dutch Research Council (14721) and an ATRAE grant from the Agencia Estatal de Investigación of the Spanish Ministry of Science, Innovation and Universities (ATR2025-164732).

## Open Practices Statement

The data, analysis scripts and materials for all experiments are available at https://osf.io/u7cx8/overview?view_only=f54e3e19d2f8479e8e10338feead572b.

## Notes

### Competing Interest Statement

The authors have declared no competing interest.

